# Ancestry Inference Using Reference Labeled Clusters of Haplotypes

**DOI:** 10.1101/2020.09.23.310698

**Authors:** Yong Wang, Shiya Song, Joshua G. Schraiber, Alisa Sedghifar, Jake K. Byrnes, David A. Turissini, Eurie L. Hong, Catherine A. Ball, Keith Noto

## Abstract

We present ARCHes, a fast and accurate haplotype-based approach for inferring an individual’s ancestry composition. Our approach works by modeling haplotype diversity from a large, admixed cohort of hundreds of thousands, then annotating those models with population information from reference panels of known ancestry. The running time of ARCHes does not depend on the size of a reference panel because training and testing are separate processes, and the inferred population-annotated haplotype models can be written to disk and reused to label large test sets in parallel (in our experiments, it averages less than one minute to assign ancestry from 32 populations to 1,001 sections of a genotype using 10 CPU). We test ARCHes on public data from the 1,000 Genomes Project and HGDP as well as simulated examples of known admixture. Our results demonstrate that ARCHes outperforms RFMix at correctly assigning both global and local ancestry at finer population scales regardless of the amount of population admixture.

**Author Summary:** Human DNA is inherited from ancestors that come from different populations across the globe and across time. Being able to identify which of those populations make up an individual’s DNA, how much they contribute, and on which chromosomes, is currently an important open research problem with many applications in the study of human diversity and history. As DNA sequencing and genotyping technology has developed, we have greater and greater amounts of data, which allows for the development of new sophisticated machine learning methods to approach this problem, and presents a need to process large amounts of data efficiently. These methods learn from examples of DNA data from known populations, and must be robust to differences in size and diversity among those reference populations. We present a new approach to this problem called ARCHes (**A**ncestry inference using **R**eference labeled **C**lusters of **H**aplotyp**es**), that models the global diversity of small segments of human DNA sequence (“haplotypes”), and the extent to which these haplotypes are associated with each of a set of population reference panels. It then computes the most likely population assignments and the points along the genome where the populations change. Our experiments show that ARCHes has superior accuracy compared to a state-of-the-art method in identifying source populations and their locations on the genome, regardless of the number of different populations present in the genome, how closely related those populations are. ARCHes is also able to model populations despite having a small amount of population reference DNA data.

## Introduction

Admixture has played an important role in shaping patterns of genetic variation among humans and other species. It is of interest at both population and individual levels and has motivated a large body of research into population demography^1, 2^ and population stratification^3^ in association studies. It has also fueled public interest in direct to consumer (DTC) services that provide estimates of ancestry proportions. In such applications, a consumer typically submits a DNA sample through a saliva collection kit and receives an individual-level report of their ancestral make-up based on genotype data.

Over the past decade, many tools have been developed to infer individual-level ancestry. One set of methods only infers global ancestry proportions, some of which model the probability of the observed genotypes using ancestry proportions and population allele frequency,^4^ while others use cluster analysis and principal component analysis (PCA).^5^ Another set of methods infer ancestral origin for genomic segments, which are then averaged over the entire genome. These methods use either SNPs (Single Nucleotide Polymorphisms) or a sequence of SNPs (*i.e.* haplotypes) as the observed variables, and estimate ancestry in each segment of the genome (called local ancestry). Compared to SNPs, haplotypes contain richer information, and can be especially powerful in differentiating geographically close populations.^6^ Among existing haplotype-based methods, both Chromopainter^6^ and HAPMIX^7^ use the Li and Stephen’s haplotype copying model,^8^ whereas RFMix^9^ uses a random forest approach, training classifiers on haplotype features in a reference panel and using a linear-chain conditional random field to model the conditional distribution of local ancestry given observed haplotypes.

As the size of public and private genotype datasets grows (*e.g.*, Ancestry has processed over 15 million human genomes), there is an increased need for methods that can efficiently and accurately perform ancestry inference on a large number of samples. Here we describe ARCHes (**A**ncestry inference using **R**eference labeled **C**lusters of **H**aplotyp**es**), a method that leverages reference panel labeled haplotype models to estimate diploid ancestry locally throughout the genome. ARCHes first uses a large set of unlabeled haplotypes to learn BEAGLE haplotype-cluster models,^10^ which are efficient at phasing and measuring haplotype frequency, for each of a number of local “windows” across the genome. These BEAGLE models are then annotated with the probability that genotype sequences from a given reference population run through a particular state. For a given test individual, ARCHes calculates the probability that the observed genotype sequence comes from a given pair of populations, followed by a genome-wide hidden Markov model to assign diploid ancestry. These trained models need only be computed once, and can be stored thereafter, allowing ARCHes to efficiently estimate the ancestry of any number of subsequent test individuals from their genotype data.

Previous studies have shown that RFMix^9^ outperforms ADMIXTURE^4^ in both global and local ancestry estimation.^11^ RFMix generally performs well at assigning ancestry at continental level but can struggle at regional level assignment, where populations may not be very differentiated. ARCHes is capable of differentiating nearby populations and performing ancestry inference at a much finer scale.

## Methods

### ARCHes Method

Our approach begins with dividing the genome into a large number of small windows (*e.g.*, 3-4 centimorgans each), such that, in a recently admixed individual, each of the maternal and paternal haplotypes in a given window are likely to each come from a single population. For each window, we construct a BEAGLE haplotype-cluster model^10^ from a large, unlabeled training set of haplotypes. A BEAGLE haplotype-cluster model is a directed acyclic graph with haplotype represented as a path traversing the graph. Each node of the graph represents a cluster of haplotypes. A BEAGLE model is often interpreted as Markov model where the states are the nodes (Figure S1), and thus as an “arbitrary order Markov model” of SNPs along a haplotype. Using a reference panel of genotypes from individuals whose ancestry is known in each window, we then annotate each state in the haplotype models with the probability that genotype sequences from a given population belong to the haplotype cluster represented by the state (Figure 1).

**Fig. 1.**
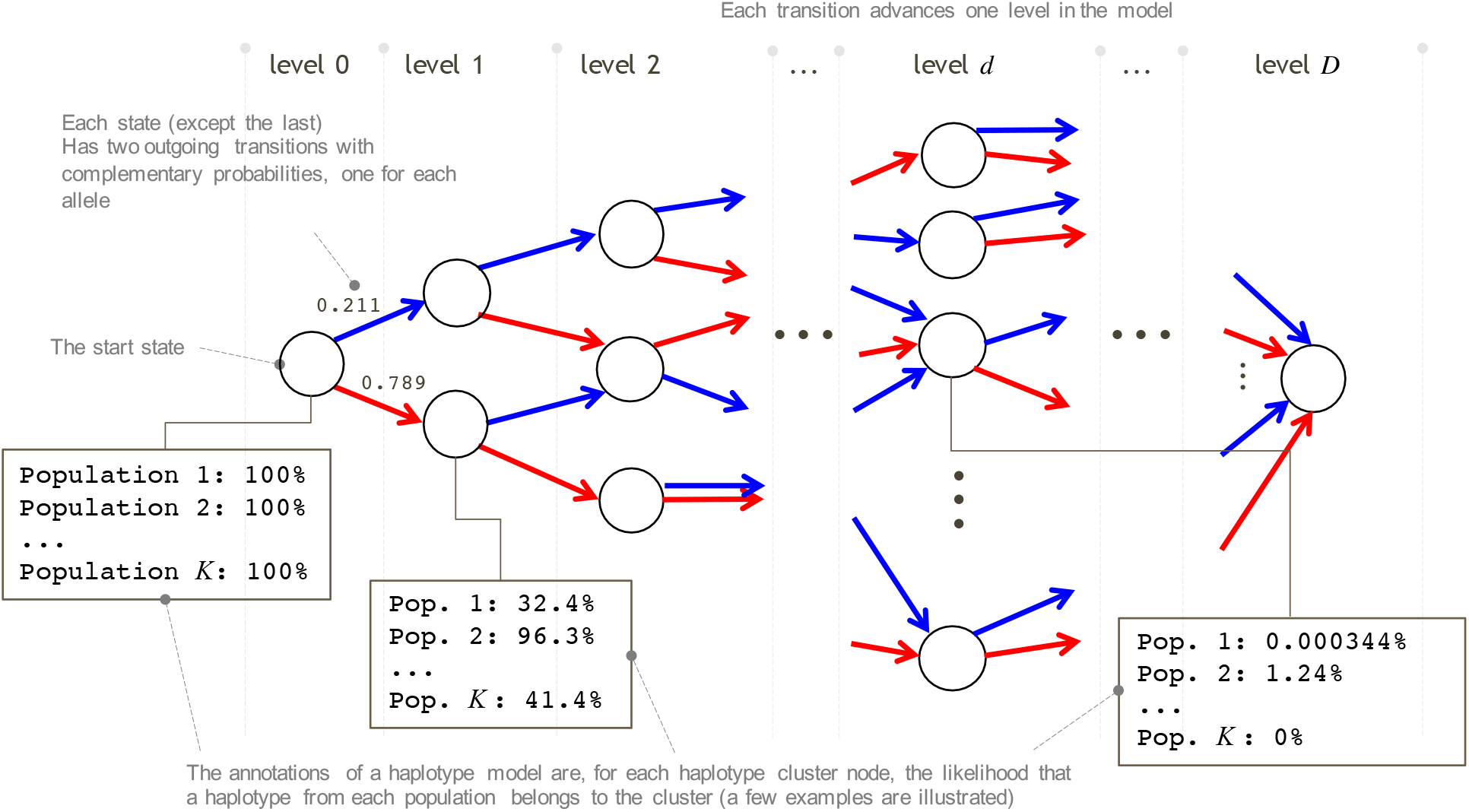
Illustration of annotating haplotype-cluster model. Each box illustrates the expected proportion of haplotypes in all the genotypes of different populations that include a certain model state at a certain level.

Given a new potentially admixed genotype sequence x, we assume that the ancestors of x are all ultimately from the *K* origin groups, and that x is admixed recently enough that relatively long haplotypes (on the scale of the genomic windows mentioned above) from each group are intact. We run a genome-wide hidden Markov model (HMM) whose hidden states are the true assignment (population label pairs) in each window. The emission probabilities are the probability distributions of diploid population assignments for each window arising from the annotated BEAGLE models and the transition probabilities (the probability that the population assignment will change at any point along the genome) are learned through an Expectation-Maximization (E-M) algorithm. We assign diploid ancestry to each window and estimate the global assignment based on the Viterbi path through this HMM. We also sample paths through the HMM to estimate the uncertainty of assignment amounts.

We describe our detailed method in the following sections, and provide pseudocode in Supplementary Materials.

### Annotating haplotype cluster models

We follow Browning and Browning^10^ in building haplotype cluster models (for practical reasons, our implementation differs in a few ways, described in Appendix S1). Briefly, we divide the genome into *W* partially overlapping windows with approximately the same number of SNPs. Within each window, we build a haplotype cluster model from a large, unlabeled set of training phased haplotypes. For simplicity, we restrict to biallelic variants, and code them as 0 and 1. Building this haplotype cluster model from a large, unlabeled set of individuals provides a “background” of haplotype diversity against which we can measure the informativeness of different haplotypes.

With a haplotype cluster model built for each window, we can then annotate populations using the haplotype cluster model. Recall that each path through a BEAGLE model corresponds to a realization of a haplotype, and each node at a given SNP represents a cluster of haplotypes that are similar near that SNP. For the genotypes of a reference individual in window *w*, x_*w*_, we compute the probability that the individual’s two haplotypes pass through two specific nodes in the era.ph *u* and at SNP *d*.

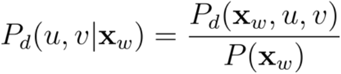

where we compute *P*_*d*_ (*u, v* | x_*w*_) and *P*(x_*w*_) using a modification of the forward-backward algorithm for hidden Markov models, treating the node as a hidden state (see Supplementary Data for pseudocode). In the following, we will refer to the HMM used to analyze the BEAGLE models as the *haplotype HMM*, and its properties as *haplotype emission probabilities* and *haplotype probabilities*. This contrasts with the *ancestry HMM* we use to smooth ancestry estimates across the genome, which is described in the subsequent section.

We then marginalize over one of the haplotypes of each diploid to create a haplotype posterior probability that the genotypes x_*w*_ in window *w* passes through node *u* at SNP *d*,

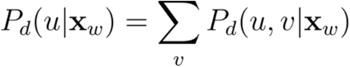

Finally, we annotate a node *u* by its average haplotype probability in a set of individuals belonging to a reference population *p,R*_*p*_ = {x_*i,p,w*,_,*i* ∈1,2,… *n*_*p*_} where *n*_*p*_ is the total number of reference samples in population. Then, we compute

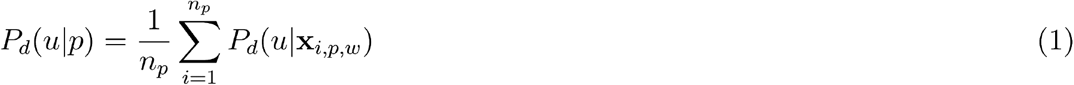

This equation gives us the probability that an individual drawn from population *P* will pass through node *u* at SNP *d* of the haplotype cluster model for window *w*.

During the annotation process, we may choose to downsample the genotypes of the reference panel by setting some genotypes at random to ‘missing’ and annotating states of the model by summing over the possible genotypes at those locations. Doing this has the effect of annotating states that represent haplotypes that are similar to those of a reference genotype, but not exactly the same, and is intended to boost performance in reference panels that have few representative examples. We may use the same reference panel individual several times in the annotation process, with a different downsampled genotype each time.

### Ancestry emission probabilities for test individuals in windows

With Equation (1) in hand, we can compute the probability that a test individual’s genotypes in a given window *w* descend from a specific pair of populations. Letting t be the unphased genotype of our test individual, we first compute the probability of t given that the two haplotypes in window *w* belong to clusters u and v of the haplotype cluster model at SNP *d*,

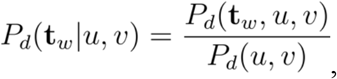

where *P*_*d*_ (t_*w*_, *u, v*) is computed using the haplotype forward-backward algorithm and *P*_*d*_ (*u, v*) is obtained by multiplying the transition matrices of the haplotype cluster model up to SNP *d* (equivalent to running the haplotype forward algorithm up to SNP *d* with all haplotype emission probabilities set equal to 1).

We then want to know the probability that the individual’s two haplotypes come from populations *P* and *q* using the information around SNP *d*. We compute this quantity by first computing the probability that a haplotype passes through nodes *u* and *v* and SNP *d* of window *w* given underlying populations *P* and *q* by averaging over the equally likely combinations of whether node *u* corresponds to population *P* and node v corresponds to population *Q* or vice versa,

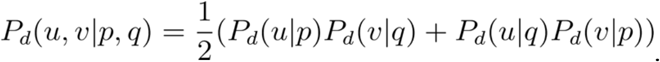

Note that this result is equivalent to assuming that the two haplotype clusters that make up a diploid sample are independent, and that the two populations that make up those haplotypes are also independent.

Now, we use the law of total probability to average over all haplotype clusters at SNP *d*, and compute the probability that the individual’s haplotype clusters at that point arise from populations *P* and *q*,

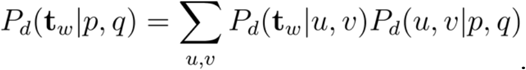

This probability weighs similarity to haplotypes in population *P* and *q* more strongly for SNPs closest to SNP *d* in window *w*; because we have no *a priori* knowledge of which part of a window is most informative about population membership, we finally compute our ancestry emission probability for a window by averaging over the population probability for every SNP in the window,

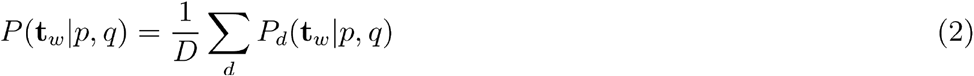

where *D* is the total number of SNPs in window *w*. This process can then be repeated for every window in the genome to obtain the probability of the test individual’s genotype in each window, given that the two haplotypes arose from any pair of populations *P* and *q*.

### Smoothing Ancestry Estimates Using a Genome-Wide Ancestry HMM

In principle, the ancestry emission probabilities computed in the previous section could be used to compute maximum likelihood estimates of diploid local ancestry in each window, one at a time. However, doing so would result in highly noisy ancestry estimates. Instead, we share information across the genome using an additional layer of smoothing via a genome-wide hidden Markov model (Figure 2). Moreover, because ancestry segments from recent admixture are expected to be longer than a single window, this model helps reduce false ancestry transitions.

**Fig 2.**
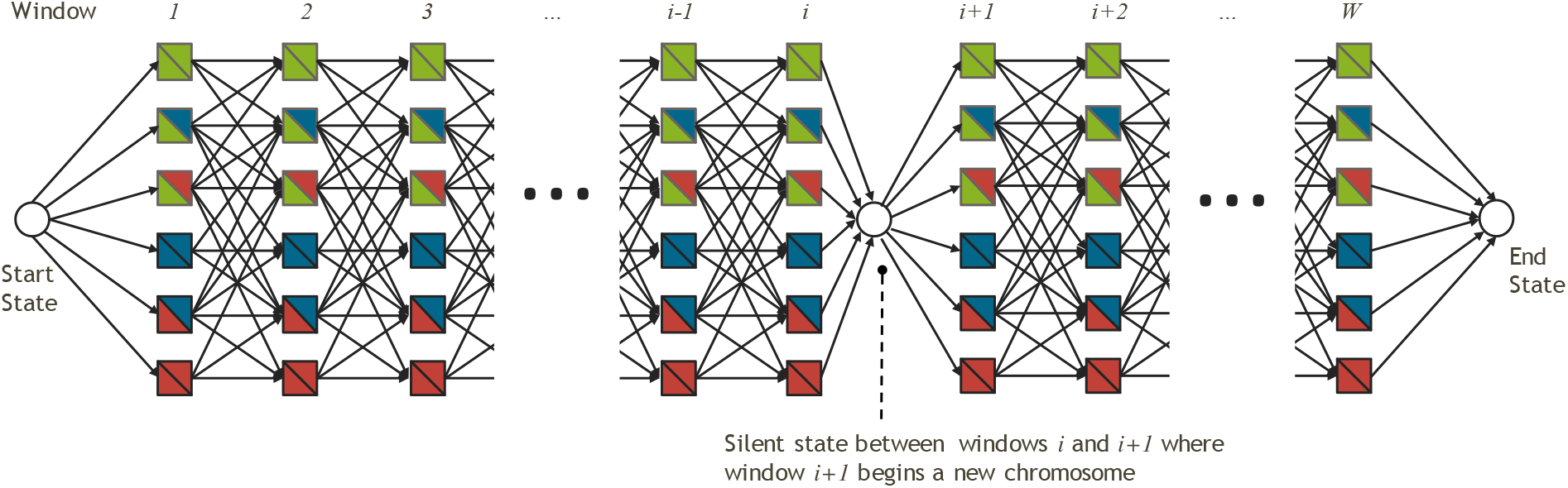
Illustration of genome wide HMM where each window has a series of emitting states, which corresponds to a population assignment (*p,q*) with *1 ≤ p ≤q ≤ K*.

If we wish to assign ancestry to *K* populations, the hidden states of our hidden Markov model are the 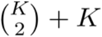 possible unphased ancestry pairs, (*p, q*), with ancestry emission probabilities window w given by equation (2). Because we model *unphased* diploid ancestry, we define a population pair as unordered, *i.e.* (*p, q*) is the same ancestry assignment as (*q, p*). Our ancestry hidden Markov model assumes that between windows ancestry can change for *one* of the two haplotypes with probability *τ*. The assumption that ancestry switches only for one of the two haplotypes within an individual is both biologically realistic (assuming individuals are admixed relatively recently) and greatly reduces the complexity of the hidden Markov model. Thus, a change occurs from (*p, q*) to (*p*′, *q*′) to any pair such that exactly one of *p*′ or *q*′ is different from *p*′ or q′. Each new ancestry pair is drawn with probability proportional to the stationary probability of that ancestry pair, *π*_*p, q*_. In full, the transition probabilities are

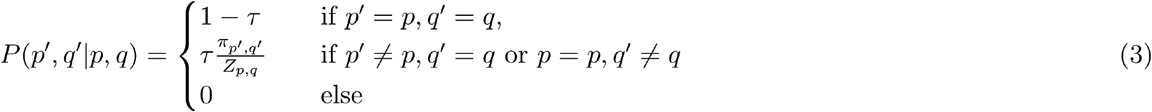

where the normalizing constant *Z*_*p,q*_ is given by summing over all accessible unphased haplotype pairs.

Between chromosomes, both ancestry pairs are allowed to change, and the ancestry at the start of each chromosome is drawn independently from that individual’s global distribution of ancestry pairs, *π*_*p,q*_ For a more formal description of how changes between chromosomes are handled, see Supplementary Data.

We initialize the *π*_*p,q*_ to a uniform distribution and *τ* to some low value, and use a modified Baum-Welch algorithm to update *π*_*p,q*_ and *τ* (see Supplementary Data). Empirically, we observed a tendency to overfit by estimating a large *τ* parameter, resulting in inference of a large number of different ancestries; thus we run a fixed number of update steps, rather than stopping at convergence.

### Estimating ancestry proportions in individuals

In principle, the value *π*_*p*_ = Σ_*q*_*π*_*p,q*_ could be used as an estimate of the admixture proportion from population *p* in an individual. However, we instead opt to use a path-based approach that also allows us to obtain credible intervals of the ancestry proportions conditioned on the inferred parameters. Specifically, we provide a point estimate of global ancestry proportions by computing the maximum probability path through the HMM using the Viterbi algorithm, and computing the proportion of windows (weighted by their length) that are assigned to population *p*. We then provide a credible interval by then sampling paths from the posterior distribution on paths, and for each one can compute the ancestry proportion in the same way as from the Viterbi path

Below we describe experiments we did for benchmarking ARCHes and RFMix^9^.

### Reference Panel and Testing Data

We build our reference panel using genotypes from proprietary candidates who explicitly provided prior consent to participate in this research project and have all family lineages tracing back to the same geographic region. All the candidates were genotyped on Ancestry’s SNP array and were analyzed through a quality control pipeline to remove samples with low genotype call rates, samples genetically related to each other, and samples who appear as outliers from their purported population of origin based on Principal Component Analysis. The reference panel contains 11,051 samples, representing ancestry from 32 global regions (Table S1). We then use 1,705 individuals from 1,000 Genomes^12^ and HGDP Project^13^ from 15 populations as testing data. We use SNP array data of individuals from the 1,000 Genomes^12^ and HGDP^13^ projects and limit them to around 300,000 SNPs that overlap with Ancestry’s SNP array. Lists of populations and associated sample counts included in reference panel and testing data are specified in Tables S1 and S2, respectively. We align populations that come from different data sources, in some cases combining populations together. For example, we combined the ancestries that are assigned to ‘England, Wales, and Northwestern Europe’ and ‘Ireland & Scotland’ to represent ancestry for ‘Britain’. We combined the ancestry that are assigned to ‘Benin & Togo’ and ‘Nigeria’ to represent ancestry for ‘Yoruba’.

### Simulation

We simulate 100 individuals with an admixture history similar to modern Latinos that admixed 12 generations ago with 45% Native American, 50% European and 5% African ancestry. We constructed 100 12-generation pedigrees and randomly selected founders from the reference panel, with the ratio of 45% Native American (from the Maya and Peru regions), 50% European (from the France, Britain, Italy, Spain and Finland regions), and 5% African ancestry (from the Yoruba region). We then simulate the DNA recombination process as above and obtained the genotypes of the descendant in each pedigree, which are admixed at roughly 45% Native American, 50% European and 5% African.

We simulate genomes of admixed individuals with ancestors from a pair of neighboring populations by simulating genotypes where 1,000 Genomes and HGDP test examples serve as the two parents, four grandparents, eight great-grandparents, or 16 great-great-grandparents of a pedigree and the admixed example evaluated is the lone descendant of that set. The examples in this test set are, on average, 50%-50% admixed, 25%-75% admixed, 12.5%-87.5% admixed, or 6.25%-93.75% admixed. We simulate 20 individuals for each of the 16 different pairings and 4 different levels of admixture, with half of them representing a minority admixture from one region, and half of them representing a minority admixture from the other region.

Since RFMix requires phased haplotypes for both query and reference individuals, we use Eagle^15^ v2 with the HRC^17^ reference panel to get phased haplotypes of the simulated individuals as well as for the individuals in the reference panel. However, ARCHes requires only the unphased, diploid genomic sequences for both query and reference individuals.

### RFMix parameters

We first used default parameters in RFMIX v2.03-r0 (https://github.com/slowkoni/rfmix). We then performed a parameter sweep using different number of generations since admixture (the -G parameter), with value of 2, 4, 6 and 8 coupled with different window sizes (set both CRF window size and random forest window size) with values of 0.2cM, 0.5cM, 100 SNPs (roughly 1cM) and 300 SNPs (roughly 3cM) on chromosome 1 of simulated pair admixed individuals. We then selected the parameters with the best performance, namely 4 generations since admixture and a window size 0.2cM, and ran RFMix on the whole genome of simulated pair admixed individuals. For simulated latino individuals, we used 12 generations since admixture and a window size 0.2 cM. For single origin individuals, we used 2 generations since admixture and a window size 0.2 cM. None of the RFMix runs used the E-M procedure or phase error correction.

### ARCHes parameters

We divide the genome into 3,882 windows of 80 SNPs each, overlapping by 5 SNPs (with some adjustments made near chromosome boundaries). We build a haplotype model for each of these windows from the phased haplotypes of 50,000 individuals that are not in the reference panel, but we tie small groups of 3-4 windows together by disallowing population assignment transitions within those groups, which allows us to set the granularity with which we assign local population assignments (there are 1,001 such window groups) and has the benefit of increased computational efficiency. ARCHes’s haplotype model annotation process is robust to missing data, which is handled by marginalizing over all possible genotypes. In fact, the annotations may benefit from intentionally downsampling reference panel genotypes so that haplotypes are considered that are similar to but not exactly the same as those in the reference panel, and the amount of downsampling and the number of downsampled genotypes used for annotation are tunable parameters of the annotation process. In our experiments, we sample each reference panel genotype sequence 100 times, each time setting 20% of genotypes to missing and annotating the 3,882 haplotype models with them. We set the initial τ_x_ parameter to be 0.01 and learned this parameter using 10 iterations of the E-M approach described above. ARCHes assigns diploid local ancestry to 1,001 windows of the genome and the global ancestry estimates are summarized from these 1,001 windows.

## Results

**Table 1.** The performance of ARCHes and RFMix on various test sets. Global concordance is the intersection between the estimated amounts for each region and the amount present in a test example, and local concordance is the number of correct assignments to each genomic window. For single-origin test examples, these measures are the same.

| Test Set Group | Test Set | Global Concordance, Average Over Test Sets<br>(Proportion of Test Sets With Superior Performance) |  | Local Concordance, Average Over Test Sets<br>(Proportion of Test Sets With Superior Performance) |  |
| --- | --- | --- | --- | --- | --- |
|  |  | ARCHes | RFMix | ARCHes | RFMix |
| 1,000 Genomes and HGDP | Single-Origin Testset Examples | **66.1%**<br>**(13/15)** | 43.5%<br>(2/15) | **66.1%**<br>**(13/15)** | 43.5%<br>(2/15) |
| Simulated Admixture | 45% Native American, 50% European, 5% African | **72.3%**<br>**(1/1)** | 65.7%<br>(0/1) | **47.8%**<br>**(1/1)** | 18.5%<br>(0/1) |
| Simulated Admixture from 16 Pairs of Neighboring Regions | 50%-50% admixed (2 simulation founders) | **60.1%**<br>**(11/16)** | 48.9%<br>(5/16) | **58.8%**<br>**(14/16)** | 41.8%<br>(2/16) |
|  | Approx. 25%-75% Admixed (4) | **63.6%**<br>**(13/16)** | 51.8%<br>(3/16) | **60.1%**<br>**(14/16)** | 44.7%<br>(2/16) |
|  | Approx. 12.5%-87.5% Admixed (8) | **65.2%**<br>**(13/16)** | 51.2%<br>(3/16) | **62.6%**<br>**(14/16)** | 46.0%<br>(2/16) |
|  | Approx. 6.25%-93.75% Admixed (16) | **66.2%**<br>**(14/16)** | 50.0%<br>(2/16) | **64.5%**<br>**(14/16)** | 47.0%<br>(2/16) |

### Accuracy for single-origin individuals

We built our reference panel using genotypes from proprietary data representing 32 population regions. We then applied ARCHes on individuals from 1,000 genomes^12^ and HGDP,^13^ representing 15 regions. (Lists of populations and associated sample sizes for both training and testing data are in Tables S1 and S2, and we describe all experimental methodology in detail, including the parameter settings for both ARCHes and RFMix in the Methods section below.) ARCHes predicts on average 66.1% of the ancestry in this test set to be from the correct region (Figure 3). The rest of the ancestry mainly came from nearby regions (Figure S2). ARCHes performs well at separating different countries within Africa, Europe, and Asia. In comparison, RFMix predicts on average 43.5% of the ancestry to be from the correct region, and the rest of the ancestry mainly came from neighboring regions, suggesting that RFMix is accurate for continental level assignments but performs less well at finer scales.

**Fig. 3.**
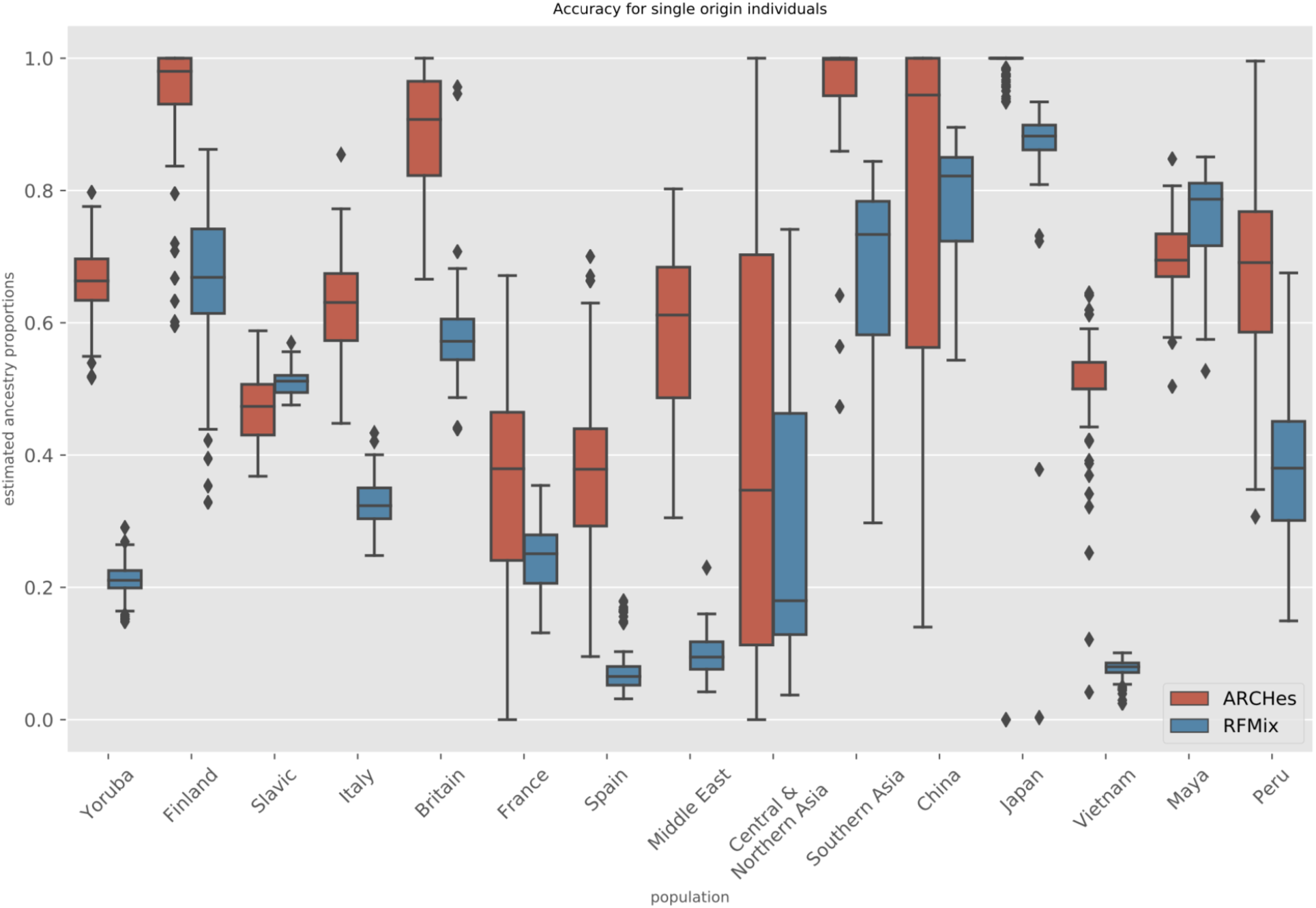
Boxplot of the estimated ancestry proportions for single-origin individuals from each testing population comparing ARCHes and RFMix.

### Accuracy for Simulated Admixed Individuals

In order to evaluate the global and local accuracy on admixed individuals, we need to know the correct ancestry throughout the genome, so we manufactured test examples from the 1,000 Genomes and HGDP data. We simulated 100 individuals using forward simulation with a pedigree mimicking Latino population history in which founders admixed 12 generations ago with 45% Native American, 50% European and 5% African ancestry. This dataset tests ARCHes’s power to differentiate continental level admixture as well as its ability to differentiate the subregions that an individual’s continental ancestry comes from.

To evaluate overall global performance on these test sets, we compute concordance as the size of the intersection between true and estimated proportions, which is the sum, for each population, of the smaller of the true global proportion and the estimated global proportion. We measured local accuracy as the proportion of genomic windows with correct diploid population assignments regardless of phase, with half credit given to a window assignment that has one population correct but the other incorrect. We find that ARCHes accurately recovers both global ancestry assignments and diploid local ancestry assignments, with average concordances of 72.3% and 47.8%, respectively (Figure S4). RFMix achieves 65.7% global ancestry concordance but failed to infer the local assignments correctly, with average diploid local ancestry concordance of 18.5%. This is due to difficulties that RFMix has in differentiatiating subregions within Europe and between Maya and Peru. The continental-level global and local concordance is 89.1% and 64.1% respectively for ARCHes, and 73.1% and 34.2% respectively for RFMix.

### Distinguishing Sub-Continental Regions

Next, we simulated genotypes for individuals with ancestry from 16 pairs of neighboring regions to test each approach’s ability to distinguish between them at global and local genomic scales Specifically, we construct test examples that are 1/2, 1/4, 1/8, or 1/16 from one region of the pair and the rest from the other region.

We measure precision and recall for each of the 11 unique regions in the set of 16 pairs (Figure 4). Precision is the amount of correctly identified ancestry divided by the amount estimated for that region and recall is the amount of correctly identified ancestry divided by the total amount of ancestry from that region that is present in the test example. ARCHes outperforms RFMix in terms of *both* precision and recall in eight of the 11 regions, and outperforms it in terms of precision in two more, and in terms of recall in one.

**Fig. 4.**
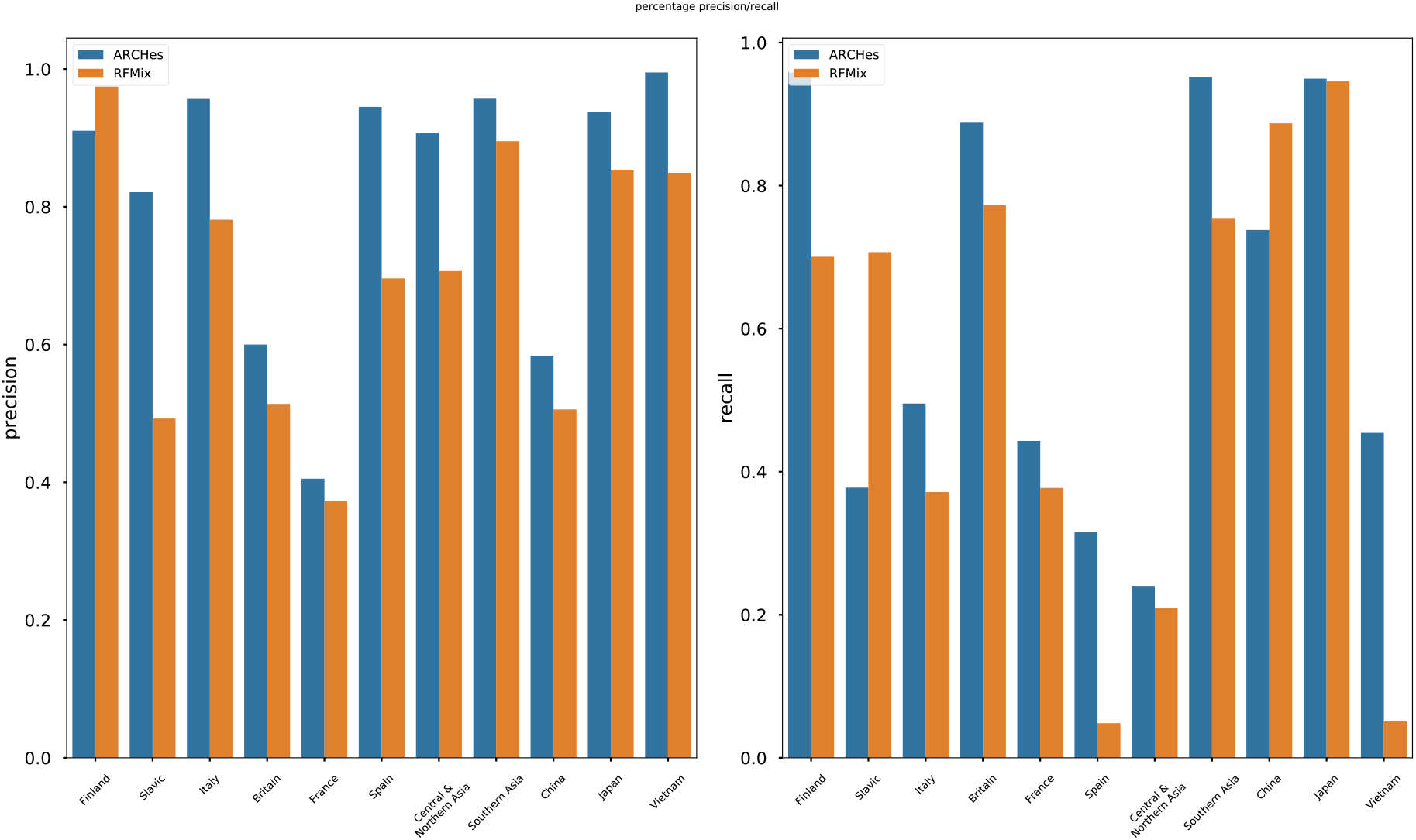
Precision/Recall for each population calculated from estimated ancestry proportions of simulated admixed individuals with ancestry from a pair of neighboring population.

Overall, ARCHes achieves more than 50% global ancestry concordance and diploid local ancestry concordance (Figure S3). There is only a small difference between global ancestry concordance and diploid local ancestry concordance on this test set, indicating that ARCHes achieves its global ancestry accuracy by estimating local ancestry accurately. It is also encouraging that ARCHes is capable of differentiating populations not only on a continental level but also on sub-continental and even country levels.

### Separate Training and Test Phases to Facilitate High-Throughput Ancestry Estimation

The ARCHes software represents a change in design that explicitly separates two phases, first model creation and annotation and second ancestry estimation, in order to make ancestry estimation both efficient and distributable. The first phase, learning the haplotype models from a large unlabeled training set and then annotating them with the reference panel populations, need only be carried out once. In order to estimate ancestry on subsequent instances, ARCHes software need only reload models and can be run on new examples at any time, distributed as necessary, and the running time depends only on the number of the number of individuals to be processed and labeled, not the size of the reference panels. In contrast, the training and testing processes of RFMix are not separate and require significantly more time per individual. We compare ARCHes’s runtime and memory usage with RFMix in Table S3.

## Discussion

Ancestry inference in large, heterogeneous sample sets is becoming increasingly important for academics, clinicians, and consumers. We developed a new approach, ARCHes, that models ancestry using rich haplotype models coupled to genome-wide information sharing. Our experiments show that ARCHes performs decisively more accurately than a state-of-the-art approach, in terms of both global and local estimation, both within and among continental scales, and among varying levels of admixture. Moreover, because our approach separates the time-consuming training step from the fast testing step, it is well-suited to apply to large scale databases.

Our approach works because haplotypes contain rich information for distinguishing subpopulations, and ARCHes’s haplotype model annotations allow it to quantitatively compare haplotypes to those of several reference panels without requiring that those reference panels be phased, contain haplotypes that are identical to that of an individual, or have similar size or diversity. Indeed, ARCHes can achieve high accuracy with reference panels containing fewer than 50 genotype examples (Figure S5). Importantly, our approach is applicable even if wholegenome single-origin training samples are not available. Because we annotate haplotype models in individual windows across the genome, we are able to utilize population-labeled partial-genome diploid or haploid genotype examples as well. That means that the accuracy of ARCHes can be improved even if a reference genotype is admixed, or if only part of it has known ancestry, by utilizing only windows of known ancestry.

Our benchmark experiments show that ARCHes is able to capture both recent and more ancient admixture by learning the genomic scale of admixture on an individual-by-individual basis: more recently admixed samples have relatively longer contiguous blocks of ancestry. This shows that ARCHes is able to be applied broadly without specific, *a priori*, parameter tuning. This feature is important for analysis of large, heterogeneous databases where it may be difficult to know the specific history of all samples involved.

ARCHes provides a fast and accurate method for inferring unphased local ancestry and combining that into estimates of diploid global ancestry. There are nonetheless several opportunities for future research. First of all, the confidence intervals provided by ARCHes are underestimated; it is possible that they can be improved by using a recalibration procedure on simulated data. Second, despite the fact that using unphased local ancestry in ARCHes helps it to overcome phasing errors, it may be desirable to provide phased local ancestry in some circumstances. Because of the modular nature of the ancestry hidden Markov model, it may be possible to extend this framework to provide phased local ancestry estimates.

## Description of Supplemental Data

Supplemental Data include five figures, three tables, and an appendix formally describing all procedures and computed values.

## Ethics Statement

All data for this research project were from subjects who provided prior informed consent to participate in AncestryDNA’s Human Diversity Project, as reviewed and approved by our external institutional review board, Advarra (formerly Quorum). All data were de-identified prior to use.

Acknowledgements

We appreciate Carlos Bustamante and Mark Koni Wright for providing RFMix software and guidance on using it.

## Supplementary Materials

**Fig. S1.**
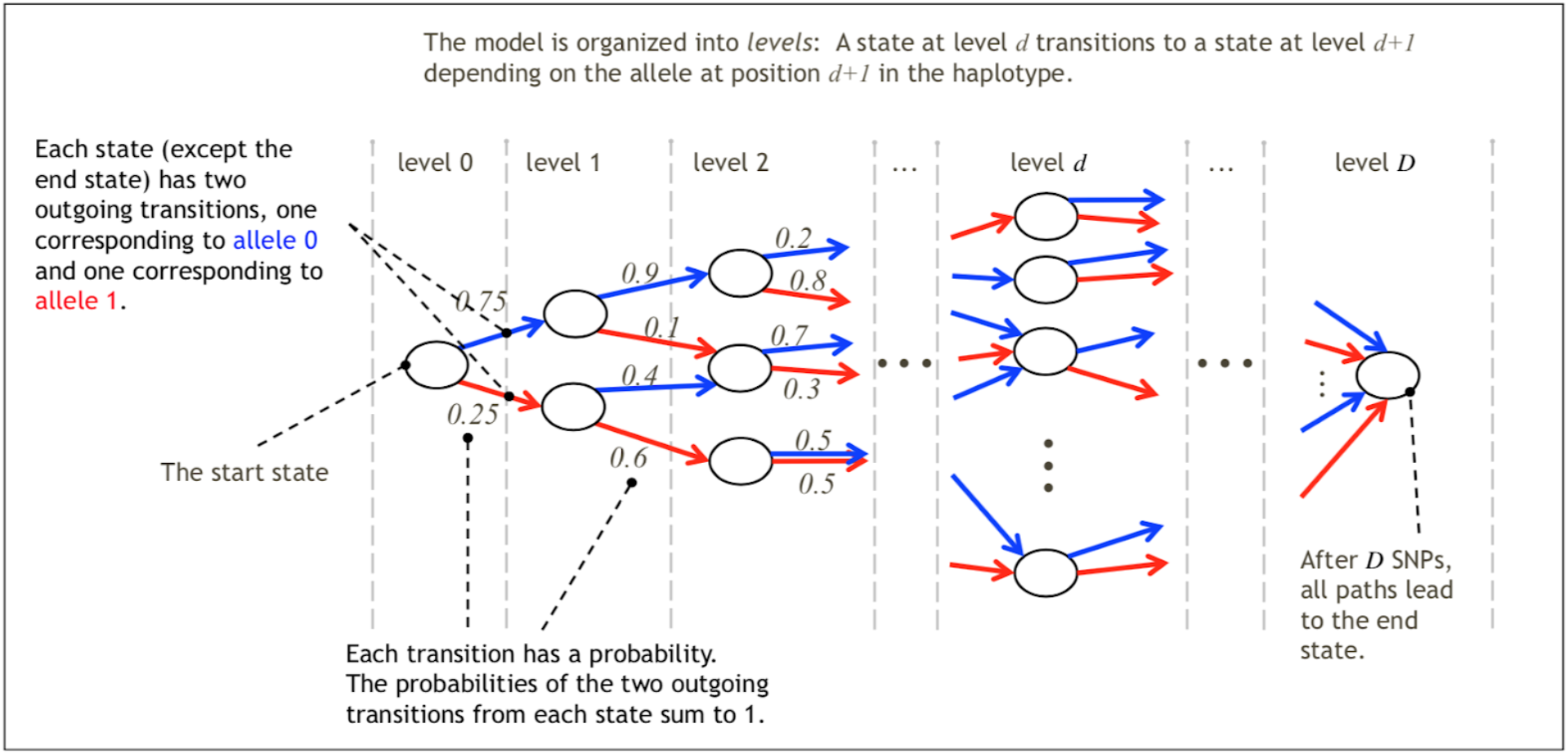
Illustration of haplotype model for one window of the genome, consisting of *D* SNPs.

**Fig. S2.**
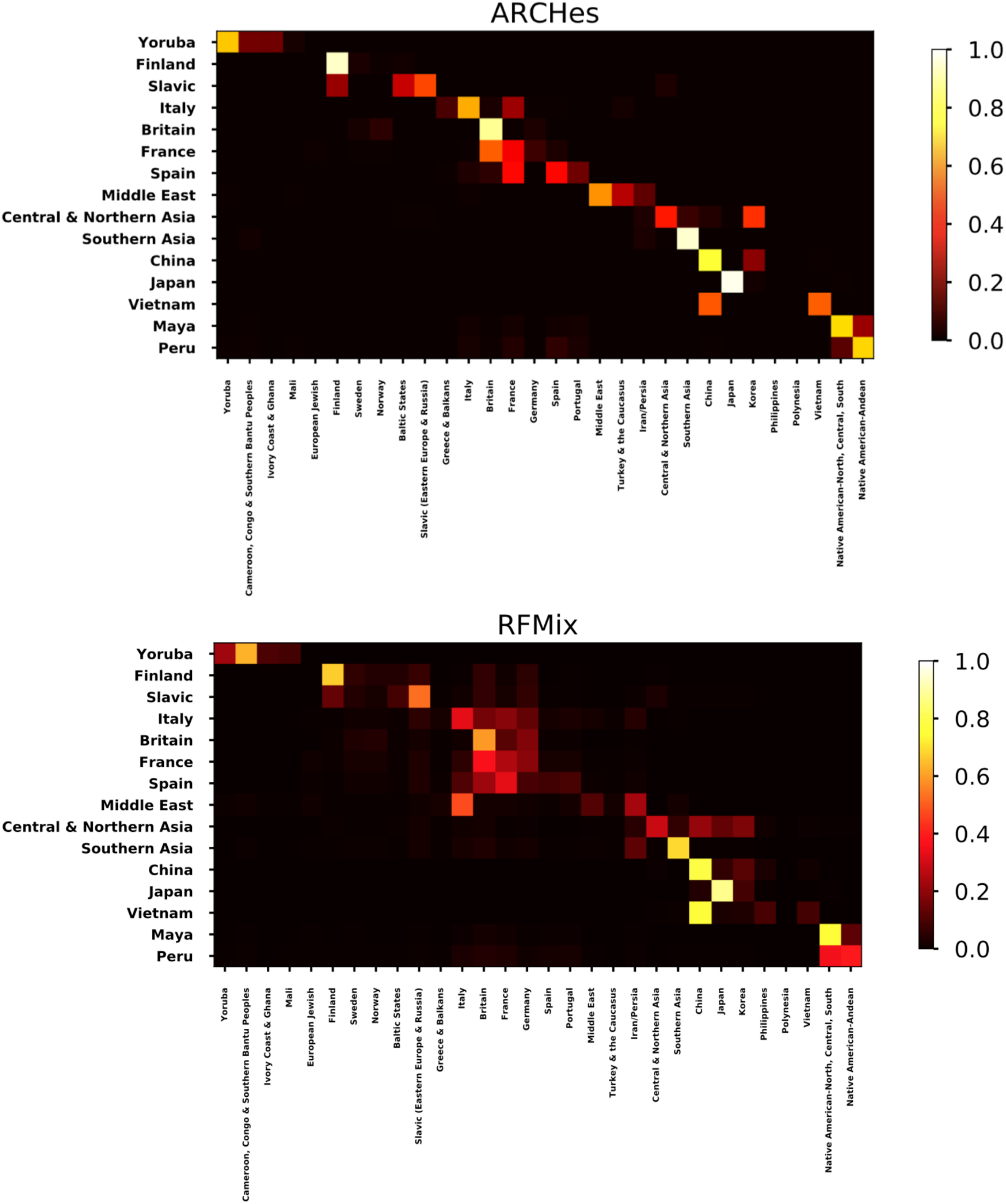
Average estimated ancestry proportions for single-origin individuals from each testing population. In this matrix figure, each row represents single-origin individuals from the testing population. Each column represents each of the possible 30 populations that the single-origin individuals might be assigned to.

**Fig. S3.**
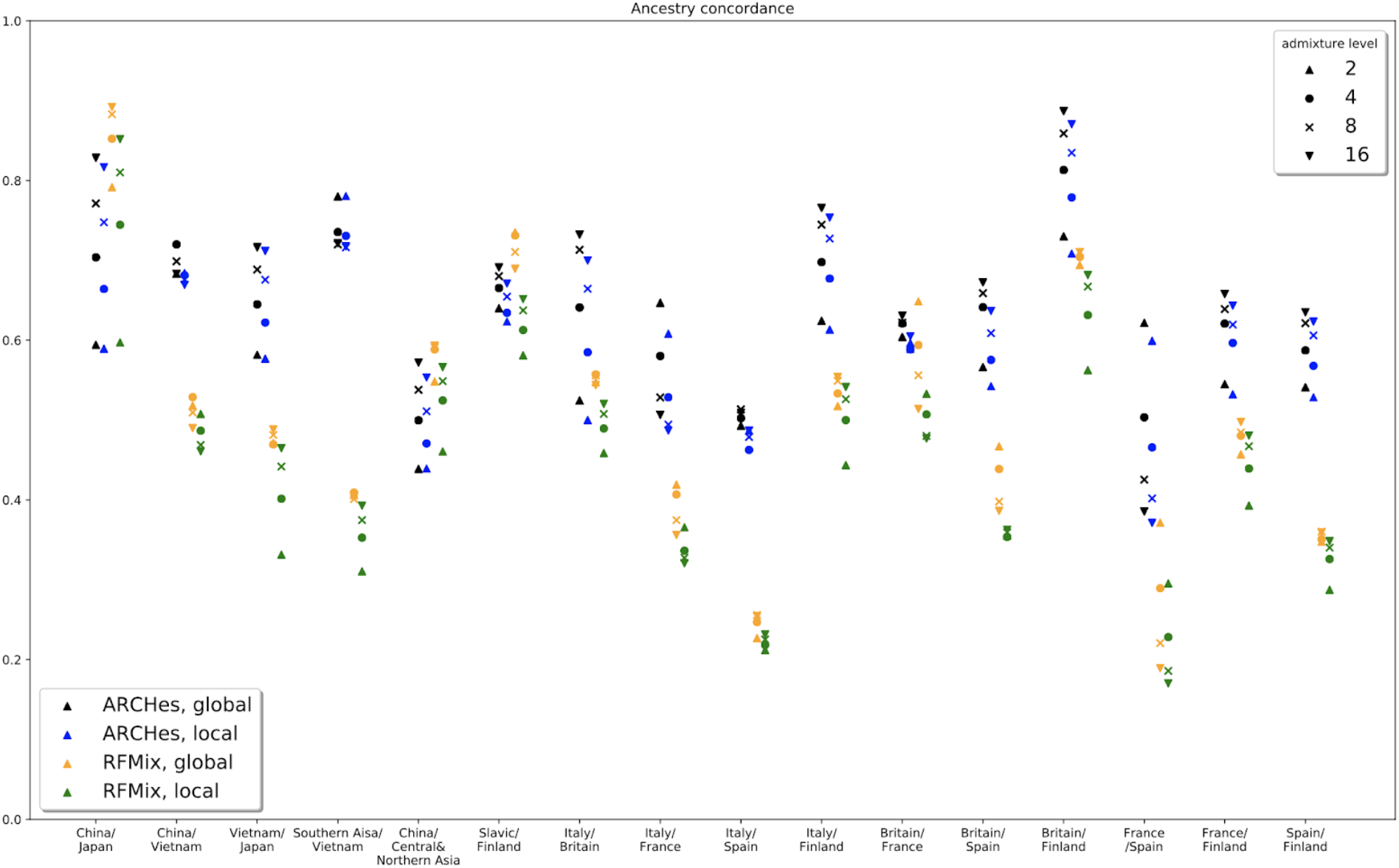
Concordance of global ancestry assignments and diploid local ancestry assignments for simulated admixed individuals from 16 different pairings of 11 populations. Admixture level is defined as x-way admixed with x founders, 1 of which belong to one population, the rest belong to another population. 2-way admixed results in 50%-50%, 4 way admixed results in roughly 25%-75%, 8 way admixed results in roughly 12.5%-87.5%, 16 way admixed results in roughly 6.25%-93.75%.

**Fig. S4.**
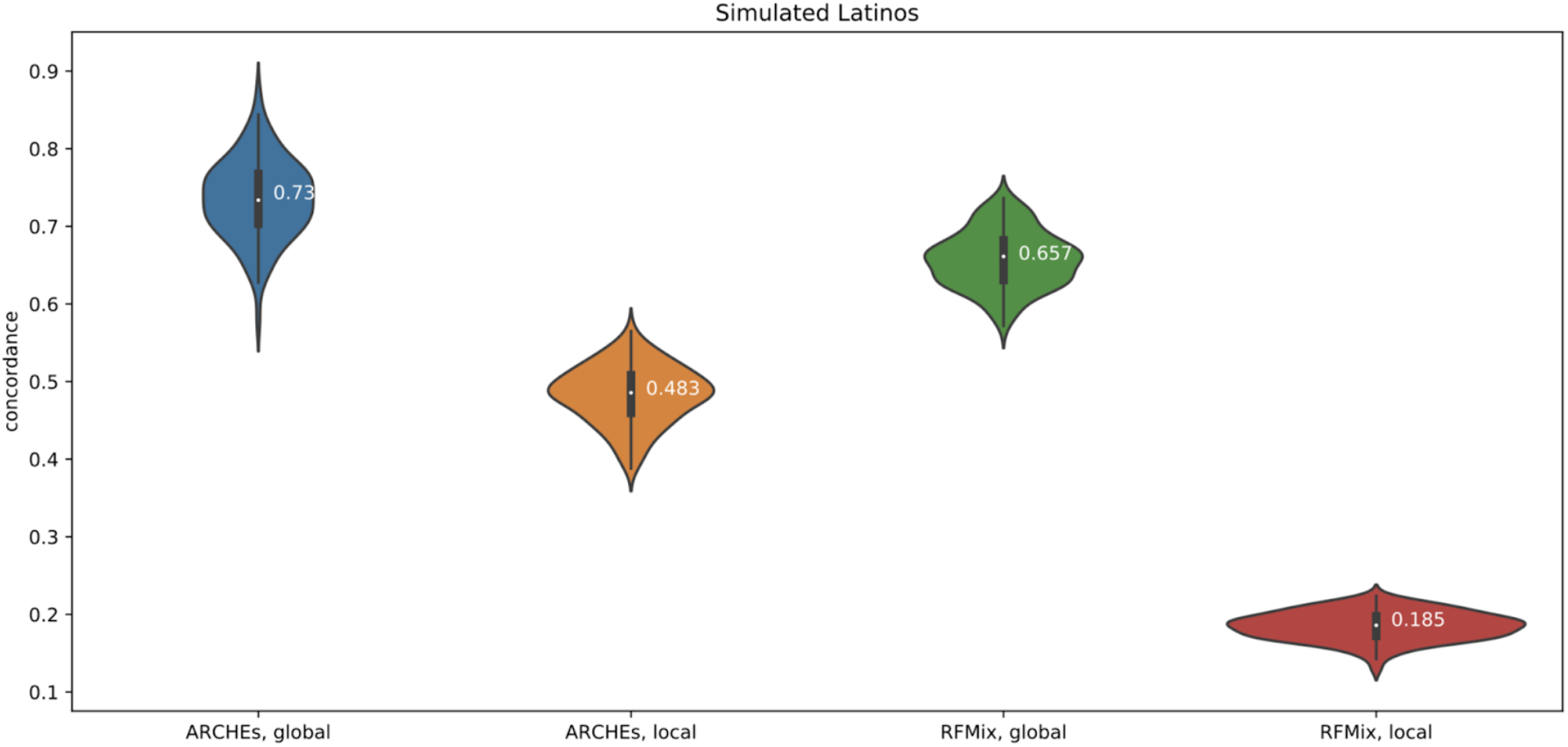
Concordance of global ancestry assignments and diploid local ancestry assignments on 100 simulated Latino individuals.

**Fig. S5.**
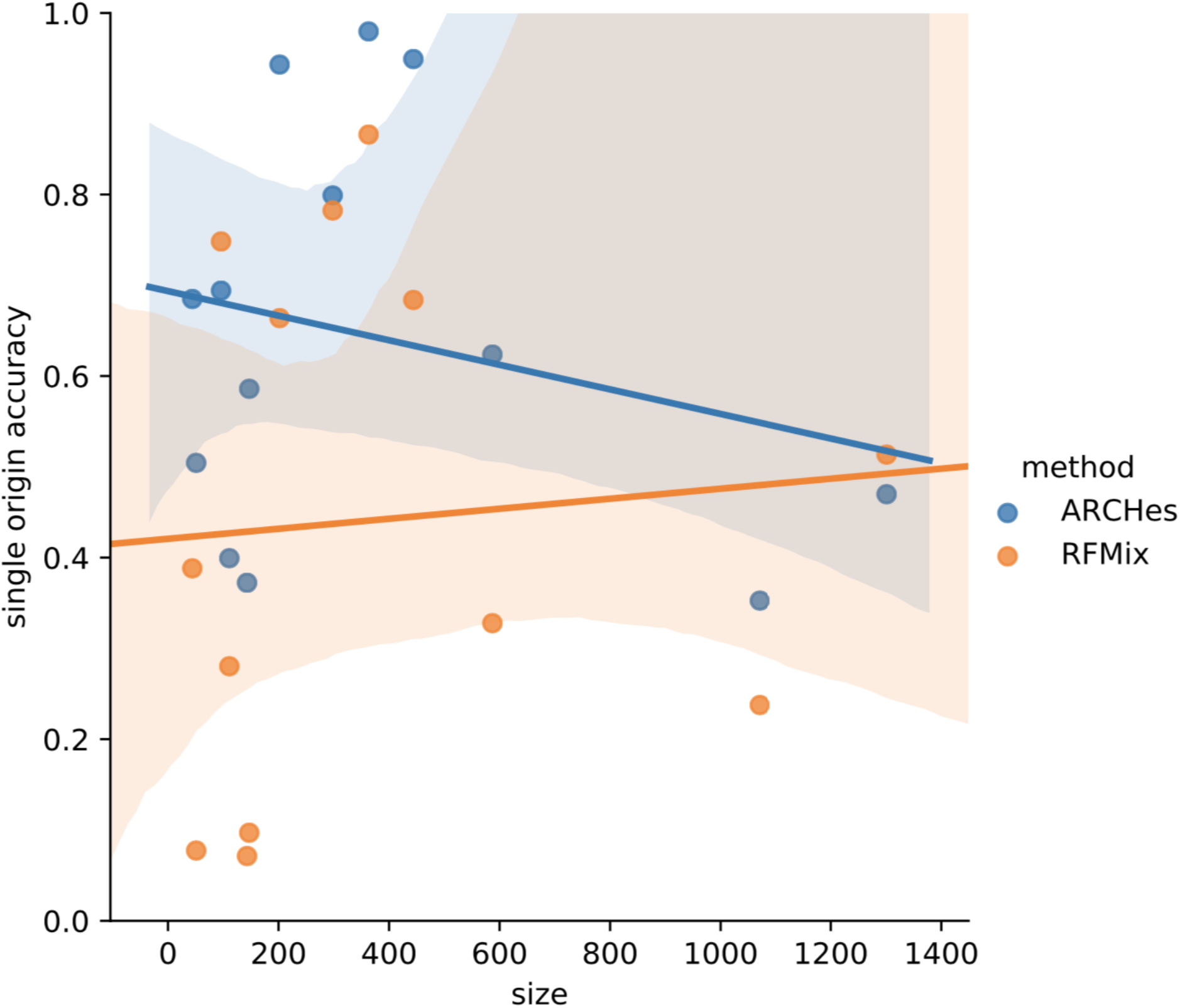
Relationship between the number of individuals in the reference panel and the accuracy for single origin individuals for each population.

**Table S1.**
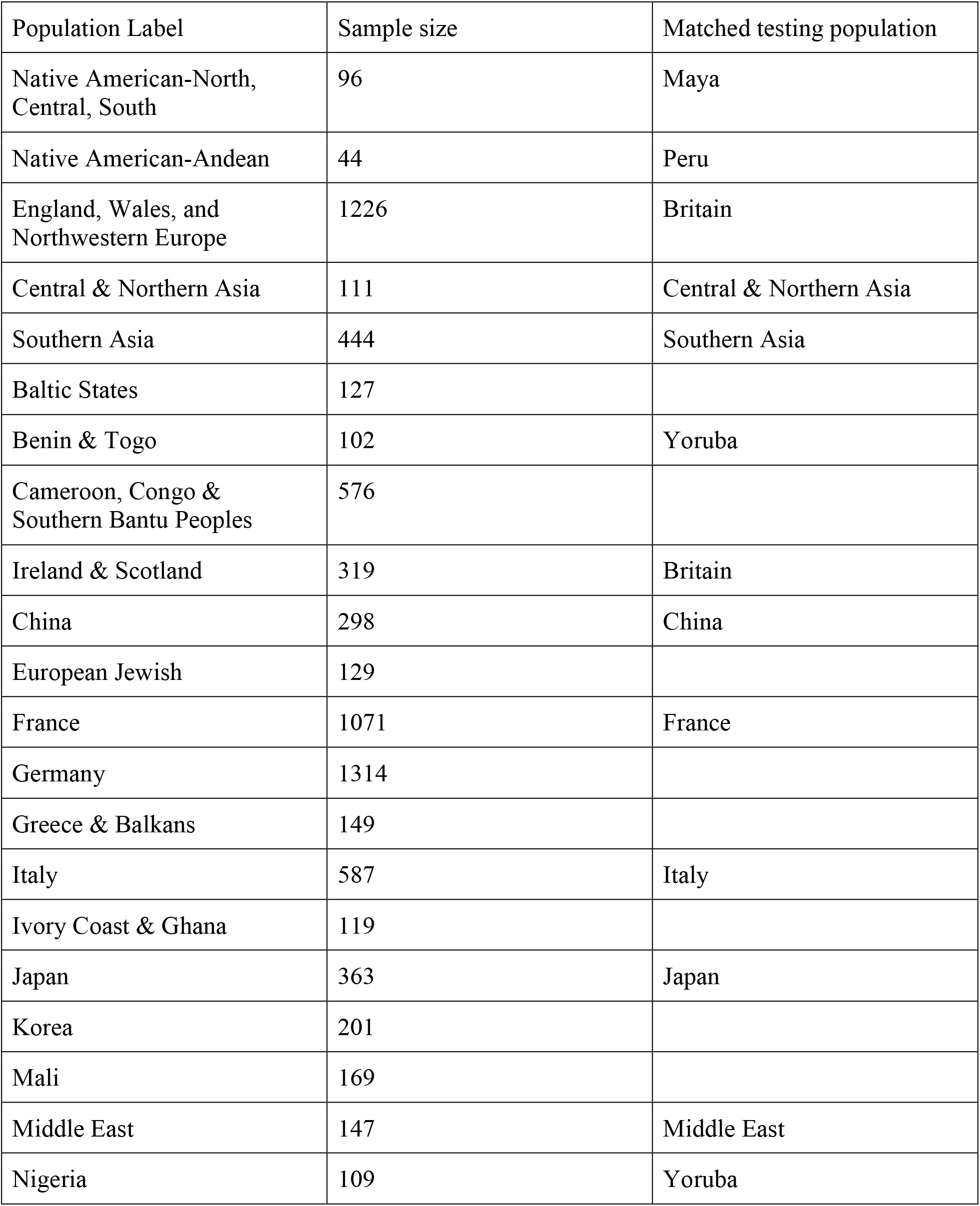
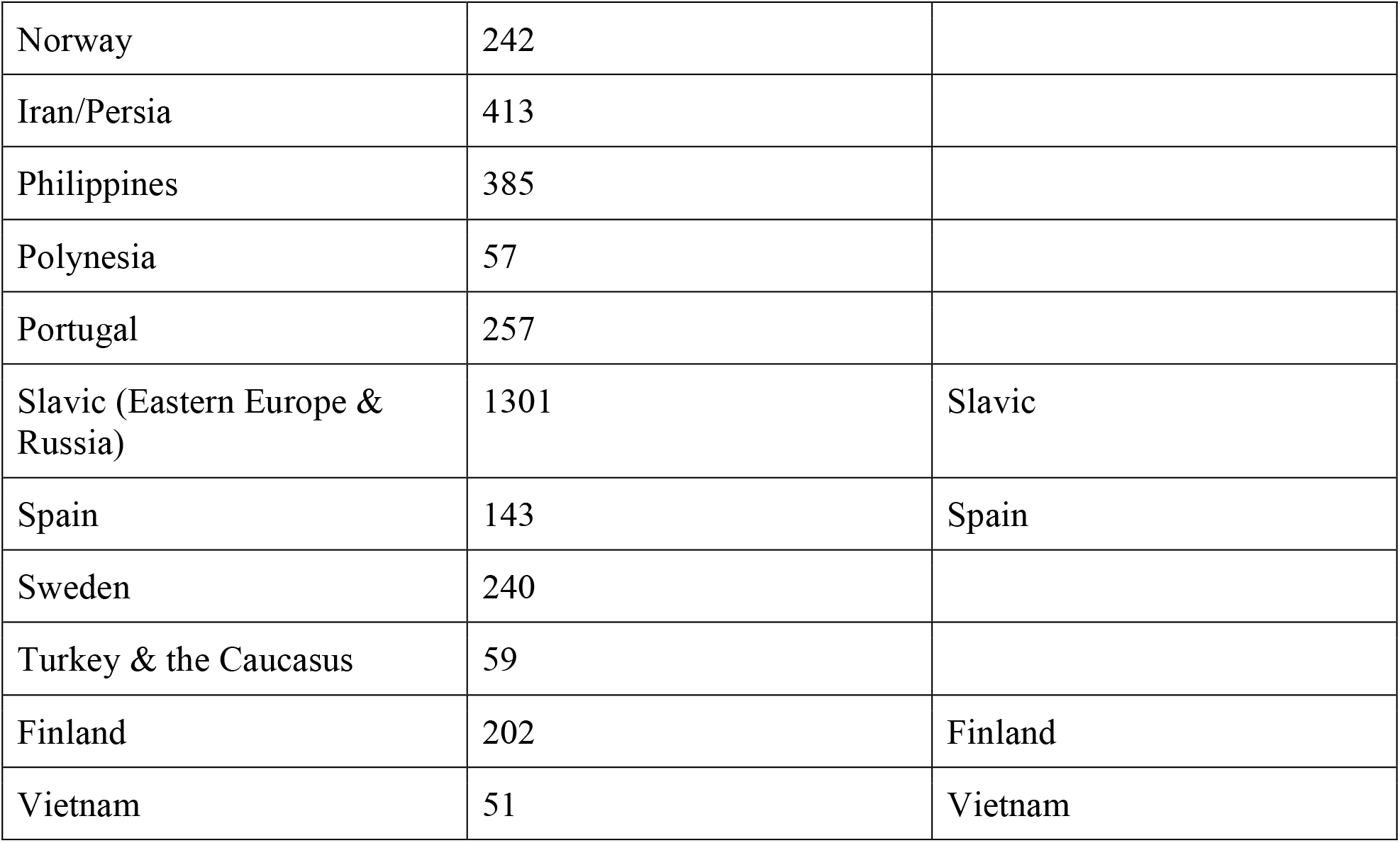
Sample size and geographic location for 32 populations in the reference panel. Some population is matched with testing population specified in Supplemental Table 2.

**Table S2.** Sample size and geographic label for testing population from HGDP and 1000 Genomes.

| Population label | Detailed label | Sample size | Source |
| --- | --- | --- | --- |
| Maya | Maya | 25 | HGDP |
| Peru | PEL(Peruvians from Lima, Peru) | 105 | 1000 Genomes |
| Central & Northern Asia | Daur, Hazara, Hezhen, Mongola, Oroqen, Tu, Uygur, Xibo, Yakut | 116 | HGDP |
| Southern Asia | Pathan, Sindhi | 48 | HGDP |
| Yoruba | YRI (Yoruba in Ibadan, Nigeria), Yoruba | 213 | 1000 Genomes, HGDP |
| China | CHS (Southern Han Chinese), Han, She, Tujia | 325 | 1000 Genomes, HGDP |
| France | French | 29 | HGDP |
| Britain | GBR (British in England and Scotland) | 104 | 1000 Genomes |
| Italy | TSI (Toscani in Italia) | 112 | 1000 Genomes |
| Japan | JPT (Japanese in Tokyo, Japan), Japanese | 134 | 1000 Genomes, HGDP |
| Middle East | Druze, Palestinian | 98 | HGDP |
| Slavic | Russian | 25 | HGDP |
| Spain | IBS (Iberian Population in Spain) | 150 | 1000 Genomes |
| Finland | FIN (Finnish in Finland) | 100 | 1000 Genomes |
| Vietnam | KHV (Kinh in Ho Chi Minh City, Vietnam) | 121 | 1000 Genomes |

**Table S3.** Run time and Memory Usage (Maximum resident set size, MaxRSS) comparison between ARCHes and RFMix. Since ARCHes trains models in a separate process, we only count the running time and MaxRSS for inferring ancestry for test individuals. However, because RFMix combines the training and testing process together, we count the running time and MaxRSS for both training and testing process for RFMix.

| Experiment | # of test individuals | Method | User time (s) | MaxRSS |
| --- | --- | --- | --- | --- |
| Single origin individual | 1705 | ARCHes | 98237 (10 CPU) | 7.9G |
|  |  | RFMix | 390443 (10 CPU) | 6.18G |
| Simulated pair admixed individual | 3200 | ARCHes | 188709 (10 CPU) | 14.8G |
|  |  | RFMix | 378838 (10 CPU) | 7.27G |
| Simulated Latino individual | 100 | ARCHes | 6814 (1 CPU) | 0.53G |
|  |  | RFMix | 389388 (10 CPU) | 8.07G |

## Appendix S1

### A Implementation of BEAGLE Haplotype Models

The haplotype models we annotate and use to compute the ancestry HMM emission probabilities are BEAGLE^10^ haplotype models, but they must be computed once and written to disk, and differ from the BEAGLE implementation in the following ways.

First, the transition probabilities are based on the haplotype counts observed in the training set, but smoothed so that all possible haplotypes have a nonzero probability. Specificially, the transition probability from a haplotype cluster corresponding to allele *a* is

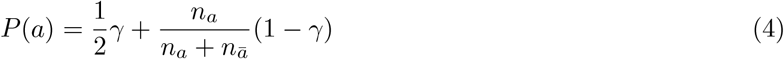

where *n*_***a***_ is the count of haplotypes in the cluster with allele *a* and *n_ā_* is the count with the alternative allele, and γ is a user-specified weight (we set γ = 10^−4^).

Second, because we build large haplotype models from hundreds of thousands of training examples, and the diploid-HMMs are quadratically larger than the haplotype models, we discard a portion of the lowest-probability states in the diploid-HMM state space after each step in the forward procedure, in order to make the procedure more efficient. Specifically, after computing the set of states (*i.e.*, possible pairs of haplotype clusters) for each SNP *d*, we sort states by forward probability and discard the least likely states, but no more than a small proportion *ϵ* (we set *ϵ* = 10^−6^) of the probability mass at level *d*.

Third, in the haplotype model building procedure, we decide when to merge two haplotype clusters based on slightly different criteria than BEAGLE. Let *n*_*x*_ and *n*_*y*_ be the respective number of haplotypes in haplotype clusters *X* and *Y*, and let *n_x_*(*h*) and *n_y_*(*h*) be the observed occurrences of a haplotype *h* in *X* and *Y*, respectively. The frequency of h in X and Y is estimated to be 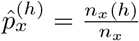 and 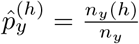, respectively. BEAGLE will not merge two clusters if

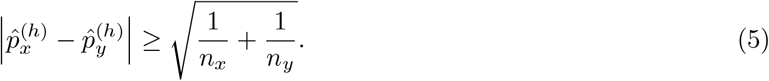

We reformulate the inequality as a hypothesis test based on Welch’s *t*-test^18^ which is, in the form of the inequalities above,

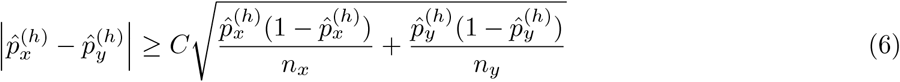

where *C* is a pre-defined constant. The concern with Welch’s *t*-test is that it is too confident in its estimation of variance when the frequency estimate is close to 0 or 1. To avoid this problem, we regularize the frequency estimate using a symmetric beta distribution as a prior. Thus, we replace the 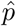 estimates with their posteriors:

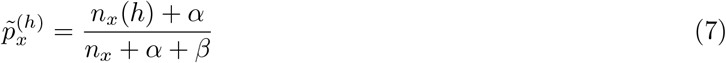

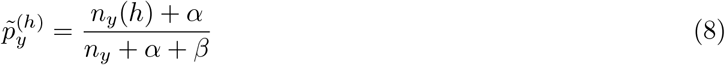

and rewrite (6) as

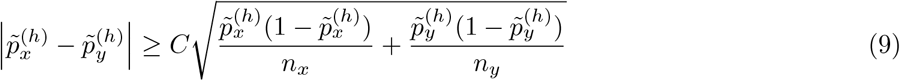

or equivalently

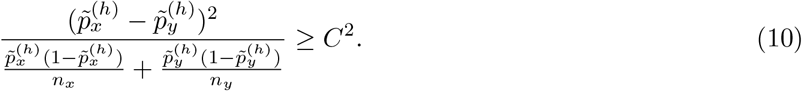

We use 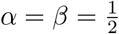 and *C*^*2*^ = 20.

### B Pseudocode for Diploid HMM Forward and Backward Procedures

#### Algorithm 1

Diploid HMM forward procedure for a sequence **x** of *D* diploid genotypes (values are all homozygous 0 or 1, heterozygous, or missing) and a model **M** of *D* + 1 levels. **M** has a start state𝕊, a transition function *t(u,a)* that maps a haplotype model state u to the state at the next level associated with the allele *a* transition (*a* ∈ 0,1), and a transition probabilty function *ρ(u,a)* that maps a haplotype model state u to the transition probability associated with allele a. The procedure populates *f*, where *f* (*u*_1_,*u*_2_) is the forward likelihood of a diploid HMM state (*u*_1_,*u*_2_). It also stores states of the diploid HMM that are consistent with the genome at each level and their outgoing transitions (and the probabilities associated with those transitions) to a data structure *a* (so that the genotype need not be re-examined during the backward procedure). The *optional* subroutine TRIM removes the diploid HMM states in a set with the lowest *f* values. It is often possible to remove a large proportion of states and yet keep (*e.g.*,) 99.9999% of the likelihood mass contained in the set of SNPs. We use TRIM only for reasons of efficiency.

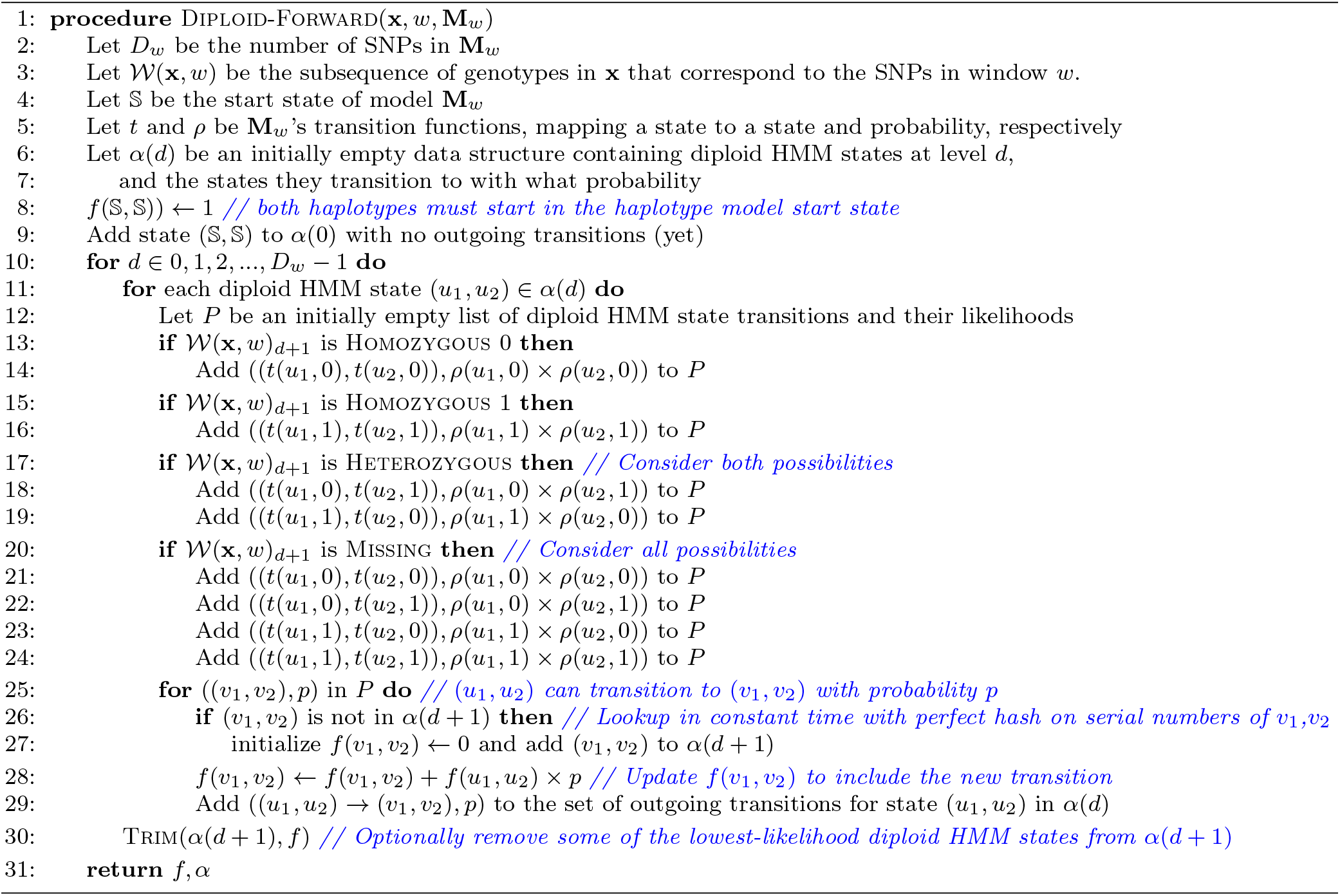

#### Algorithm 2

Diploid HMM backward procedure (see Dlploid-Forward). The procedure populates *b*, where *b*(*u*_1_,*u*_2_) is the backward likelihood of a diploid HMM state (*u*_1_,*u*_2_). *D* is the number of SNPs in the window associated with the haplotype model, *α* is the set of diploid HMM states at each level and their probabilistic outgoing transitions as computed by diploid-forward.

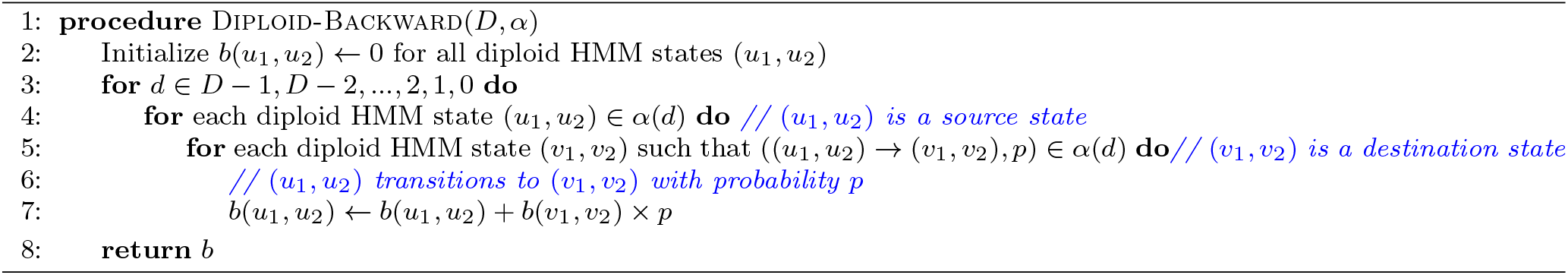

#### Algorithm 3

Diploid HMM forward-backward procedure (see Diploid-Forward and Diploid-Backward). The procedure populates *f* and *b*, where *f*(*u* sub>1, *u* sub>2) is the (“forward”) likelihood that a path through the diploid HMM ends in state (*u*_1_, *u*_2_) after emitting *d* alleles (where *d* is the level of *u*_1_ and *u*_2_) of a haplotype in the input genotype sequence **x**, and b(u1, u2) is the likelihood of all paths from (*u*_1_, *u*_2_) to the end state. The probability *P*_*d*_(*u*_1_, *u*_2_|x) that the haplotypes of genotype sequence **x** belongs to clusters *u*_1_ and *u*_2_ is calculated as 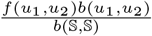, where 𝕊 is the start state of model **M** and *f* and *b* are computed by this procedure.

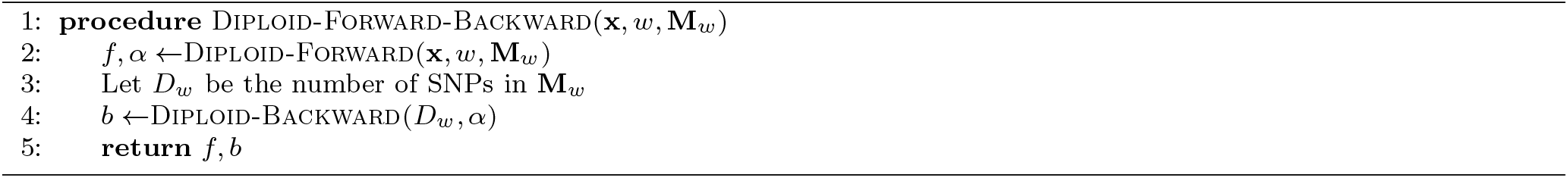

### C Computing Forward-Backward on the Genome-Wide Ancestry HMM, and Updating π and *τ* Transition Probability Parameters

The genome-wide ancestry HMM computes the likelihoods that a test instance’s genotype sequence, **t**, in a genomic window *w* (denoted **t**_*w*_) is explained by populations *p* and *q* for a set of populations and genomic windows. It is parameterized by π_**t**_ and *τ*_***t***_,^1^ which are typically learned for a specific test instance. The ancestry HMM representing a *K* populations and a set of SNPs on multiple chromosomes has a single silent (non-emitting) state before the first, after the last, and in-between each chromosome, and a series of 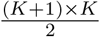 emitting states for each window of each chromosome, each corresponding to a population assignment (*p, q*) with 1 ≤ *p* ≤ *q* ≤ *K*. Figure 4 illustrates such a genome-wide HMM with *K* = 3 populations. Let the emitting state corresponding to window *w* and population assignment (*p, q*) be denoted *S*_*w*_,_*p*_,_*q*_. Its emission probability *P*(**t**_*w*_ |*p, q*) is precomputed and fixed based on the genotype **t**_*w*_ in window *w*. Let *S*_*c*_ represent the silent state that preceeds the emitting states correpsonding to windows on chromosome c. Thus the start state of the HMM is *S*_1_ (and if the HMM represents C chromosomes, the end state would be *SC*+1). Let 𝒞(*c*) map a chromosome number to the window that begins the chromosome. Then, our HMM transitions from silent states to emitting states *S*_*c*_ → *S*_*𝒞 (****c***)_,_*p*_,_*q*_, from emitting states in the last window of a chromosome to a silent state *S*_*𝒞* (c+1)_1,*p*_,_*q*_ → *S*_*c*_+1, and from emitting states to emitting states for windows w that are not the first or last in a chromosome *S*_*w,p,q*_. → *S*_*w*+1,*P*′_,_*q*′_. A transition from *S*_*w,p,q*_ → *S*_*w*+1,*P*′_,_*q*′_ represents a change in population assignment between windows *w* and *w* + 1 if *p* ≠ *p*′or q ≠ *q*′

The transition probabilities from a silent state to an emitting state *S*_*c*_ → *S*_*𝒞* (c), p,q_ is π_**t**_,(_*p*_,_*q*_), where π_**t**_ is a learned parameter vector over all possible assignments (*p, q*) (1 ≤ *p* ≤ *q* ≤ K) indicating a global assigment preference. The transition probability from a state in the last window of a chromosome to a silent state *S*_*𝒞*_(_*c*_)-1,_*p*_,_*q*_ → *S*_*c*_ is always 1, and transitions between emitting states on the same chromosome, from state (*p, q*) in window *w* to state (*p*′, *q*′) (with *p*′ ≤ *q*′) in window *w* + 1, are as follows:

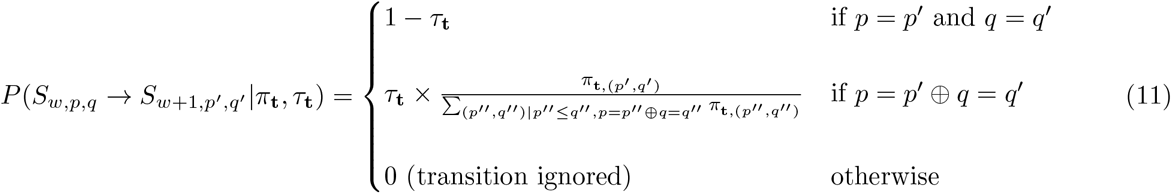

where *τ*_**t**_ is a parameter representing the probability of changing population assignment that enforces the bias against changing population assignments from window to window (⊕ is the exclusive or operator). We initialize *π*_**t**_ to a uniform distribution, and *τ*_**t**_ to a (typically low) initial value and learn *π*_**t**_ and *τ*_**t**_ using expectation-maximization over a number of iterations (similar to the standard Baum-Welch algorithm^16^ except that *π*_**t**_ and *τ*_**t**_ are tied to all state transition probabilities).

Let *F*_**t**_(*s*) be the forward probability, the sum probability of all paths through the Ancestry HMM (as opposed to the *haplotype* HMM used to calculate per-window emission probabilities) that start in the start state and end in state *s* (including the emission of state *s*) and *B*_**t**_(*s*) be the backward probability of all paths through the HMM that start in state *s* (excluding emission) and end in the end state. *F* and *B* are computed recursively as follows.

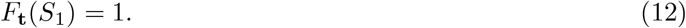

For the emitting states in the first window of a chromosome,

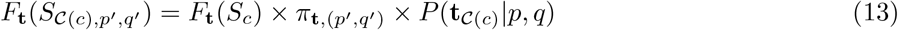

for all *p*′ and *q*′. When a window *w* is not the first window of a chromosome,

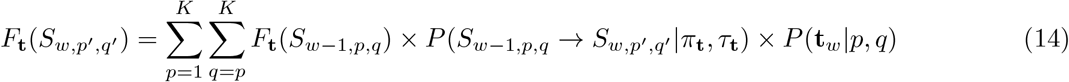

where 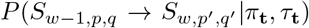 is given by (11). The forward probability of the silent state preceding chromosome *c* is

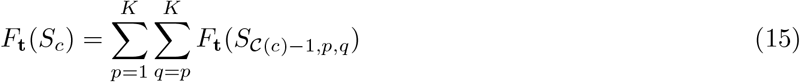

Similarly, if there are *C* chromosomes in the model,

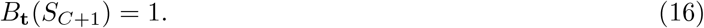

For the last window on chromosome *c*,

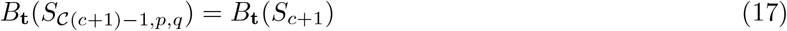

for all *p* and *q*. When window *w* is not the last window on a chromosome,

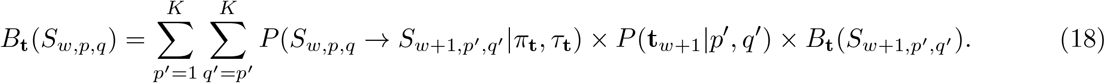

Finally, for the silent state preceding chromosome *c*,

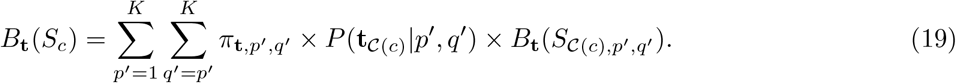

After computing Ft and Bt, we compute the expectation for each π_**t**,(p,q)_ as

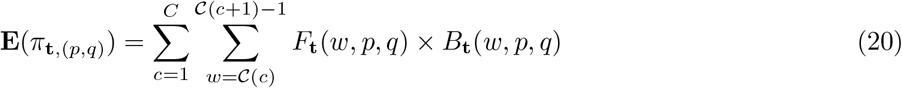

and reset each π_**t**,(p,q)_ to the value that maximizes the likelihood of E(π_**t**,(p,q)_):

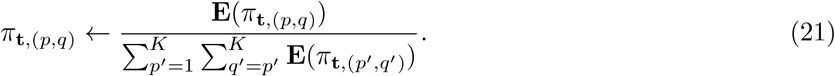

We learn π_**t**_ in a similar fashion, by updating it based on the expected number of transitions that do not change assignment, compared to all transitions. If there are C chromosomes,

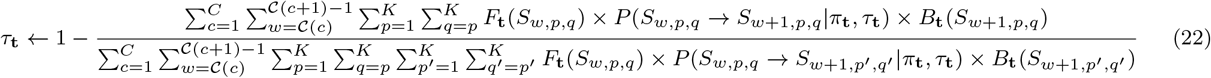

### D Computing the Viterbi Path

The Viterbi path is the single most likely path (relative to a genotype sequence t) through the genome-wide HMM **V**_**t**_ = ⟨ V_t,1_, V_t,2_,…, V_t,w_ ⟩, where each V_t,w_ is an assignment (p, q) in a window w, 1 ≤ w ≤ W.

To compute **V**, we must first define M_**t**_, where *M*_***t***_(s) is the probability of the most likely path through the HMM that start in the start state and end in state s (including the emission of state s), analagous to the forward probability *F*_**t**_(s) AppendixC, but referring to the probability of the single most likely path instead of the sum probability of all paths.

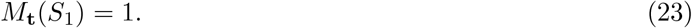

For the emitting states in the first window of a chromosome,

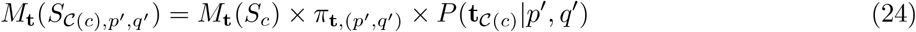

for all p′ and q′. When a window *w* is not the first window of a chromosome,

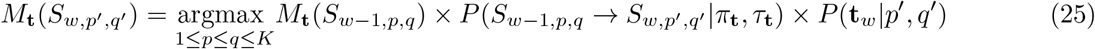

And for a silent state that is not the start state,

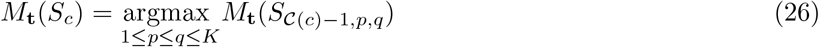

The Viterbi path **V** is then defined for windows that are the last window in a chromosome, *c*, as

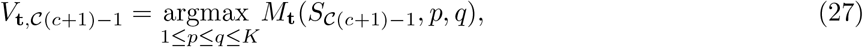

and for all other windows as

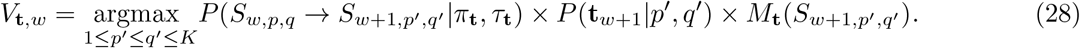

### E Computing Path Samples

Let choose be a operator that chooses an argument with a probability relative to an expression so that 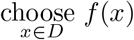 returns x with probability 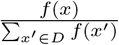 Then a stochastic path **Q** for a genomic sequence **t** is defined over all windows 1 ≤ w ≤ W as follows. For windows that are last in a chromosome, *c*,

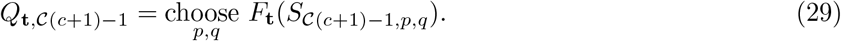

For other windows *w*,

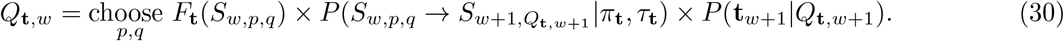

## Footnotes

1 The subscript **t** may be dropped from these and other terms when there is only one test genotype instance in question.

